# CELF1 is an EIF4E binding protein that promotes translation of epithelial-mesenchymal transition effector mRNAs

**DOI:** 10.1101/640300

**Authors:** Arindam Chaudhury, Rituraj Pal, Natee Kongchan, Na Zhao, Yingmin Zhu, Emuejevoke Olokpa, Shebna A. Cheema, Sonia del Rincon, Lucas C. Reineke, Sufeng Mao, Richard E. Lloyd, Marco Sardiello, Jeffrey M. Rosen, Joel R. Neilson

**Affiliations:** Department of Molecular Physiology and Biophysics, Baylor College of Medicine, Houston, TX 77030, USA; Dan L. Duncan Comprehensive Cancer Center, Baylor College of Medicine, Houston, TX 77030, USA; Department of Molecular and Human Genetics, Baylor College of Medicine, Houston, TX 77030, USA; Department of Molecular and Cellular Biology, Baylor College of Medicine, Houston, TX 77030, USA; Lester and Sue Smith Breast Center, Baylor College of Medicine, Houston, Texas 77030, USA; Protein and Monoclonal Antibody Production Core, Baylor College of Medicine, Houston, TX 77030; Gerald Bronfman Department of Oncology, McGill University, Lady Davis Institute for Medical Research, Segal Cancer Centre of the Jewish General Hospital, Montreal, Quebec, H3T 1E2, Canada; Department of Virology and Microbiology, Baylor College of Medicine, Houston, TX 77030, USA

## Abstract

Mounting evidence is revealing a granularity within gene regulation that occurs at the level of mRNA translation. Within mammalian cells, canonical cap-dependent mRNA translation is dependent upon the interaction between the m^7^G cap-binding protein eukaryotic initiation factor 4E (eIF4E) and the scaffolding protein eukaryotic initiation factor 4G (eIF4G), the latter of which facilitates pre-translation initiation complex assembly, mRNA circularization, and ultimately ribosomal scanning. In breast epithelial cells, we previously demonstrated that the CELF1 RNA-binding protein promotes the translation of epithelial to mesenchymal transition (EMT) effector mRNAs containing GU-rich elements (GREs) within their 3’ untranslated regions (UTRs). Here we show that within this context, CELF1 directly binds to both the eIF4E cap-binding protein and Poly(A) binding protein (PABP), promoting translation of GRE-containing mRNAs in mesenchymal cells. Disruption of this CELF1/eIF4E interaction inhibits both EMT induction and experimental metastasis. Our findings illustrate a novel way in which non-canonical mechanisms of translation initiation underlie transitional cellular states within the context of development or human disease.

## Introduction

Emerging evidence suggests that regulatory control at the level of mRNA translation contributes significantly to gene expression and function(1–4). Consistent with this evidence, recent findings indicate that coordinated changes in post-transcriptional regulatory networks can alter cellular phenotype and behavior (2,5). A core cellular process underlying development, tumor metastasis, and tumor radiation and chemoresistance is epithelial to mesenchymal transition (EMT) (6,7). Since the metastasis and resistance to insult associated with cellular de-differentiation are the foremost cause of cancer lethality (8,9), it is critical to develop a better understanding of the mechanisms by which EMT, and thus these two characteristics, are promoted. We previously defined a translational regulatory circuit driving EMT *in vitro*, and cancer progression *in vivo*, in breast epithelial cells (10). The key regulator of this circuit is the RNA binding protein CELF1, which promotes translation of EMT effector mRNAs by binding to <u>G</u>U-rich <u>e</u>lements (GREs) within the 3’ untranslated regions (UTRs) of these mRNAs.

Canonical eukaryotic translation initiation is a well-defined and ordered process that guides the assembly of ribosomes onto m^7^G-capped mRNAs (11). The m^7^G cap structure on the mRNA is first bound by the eIF4E protein. eIF4E recruits the other two members of the tripartite eIF4F complex - the scaffold protein eIF4G1 and the RNA helicase eIF4A - to the cap structure (11). eIF4G1 bridges eIF4E and eIF4A to Poly-A binding protein (PABP) and the eIF3-eIF1-eIF1A complex, the latter of which guides the (small) 40S ribosomal subunit, bound with methionine-charged initiator transfer RNA and eIF2 to form the 48S pre-initiation complex (PIC) (12). EIF4G1’s role is so central to cap-dependent translational initiation that cleavage of this protein in virally infected (13,14) cells is sufficient to broadly disrupt cap-dependent translation, giving the virus complete control of the cell.

It has now been well-documented that both oncogenes and signaling pathways regulating expression of these oncogenes together converge on the regulation of mRNA translation (2,15). Indeed, within the process of cellular transformation, mechanisms at each stage of translation and the ribosomal machinery may be co-opted to fostering translation that perpetuates the oncogenic program (16–18). This motivated us to more closely examine how CELF1 regulates translation of GRE-containing EMT effector mRNAs during EMT of breast epithelial cells.

## Results

### CELF1 interacts directly with eIF4E and PABP at the m^7^G cap, independent of intact eIF4G1

Our previous work suggested that CELF1’s control of GU-rich element-containing EMT effector mRNA translation is mediated at the level of 5’ m^7^G cap-dependent translation initiation (10). We first confirmed and extended these findings by examining an additional CELF1 regulatory target within this context. We fused the 3’ UTRs of a subset of GRE-containing mRNAs (*JUNB*, *CRLF1*, *SNAI1*, and *SSBP2*) downstream of the *Renilla* luciferase coding sequence in the *pRL-TK-CXCR4-6x* (19) reporter plasmid (Figure 1a). Transfection of the GRE-containing 3’ UTR reporters into MCF10A cells, either untreated or treated with TGF-β to induce EMT, revealed a significant increase in reporter activity specific to treated cells Figure 1a), independent of any differences in relative mRNA expression (Figure 1b). In parallel, we built a battery of bicistronic constructs in which the same thymidine kinase promoter was utilized to drive expression of the firefly luciferase coding sequence, followed by the encephalomyocarditis virus (EMCV) internal ribosome entry site (IRES), the *Renilla* luciferase open reading frame and individual GRE-containing 3’ UTRs (*pFR-EMCV*) (Figure 1c). In contrast to cap-dependent *Renilla* luciferase expression, no differences in relative reporter expression driven by the IRES were observed in a comparison of the epithelial and mesenchymal states (Figure 1c, 1d). These results confirmed and extended our previous observation that translational control of mRNAs harboring GREs within their 3’ UTRs during EMT is mediated at the level of 5’ m^7^G cap-dependent translational initiation (10).

**Figure 1.**
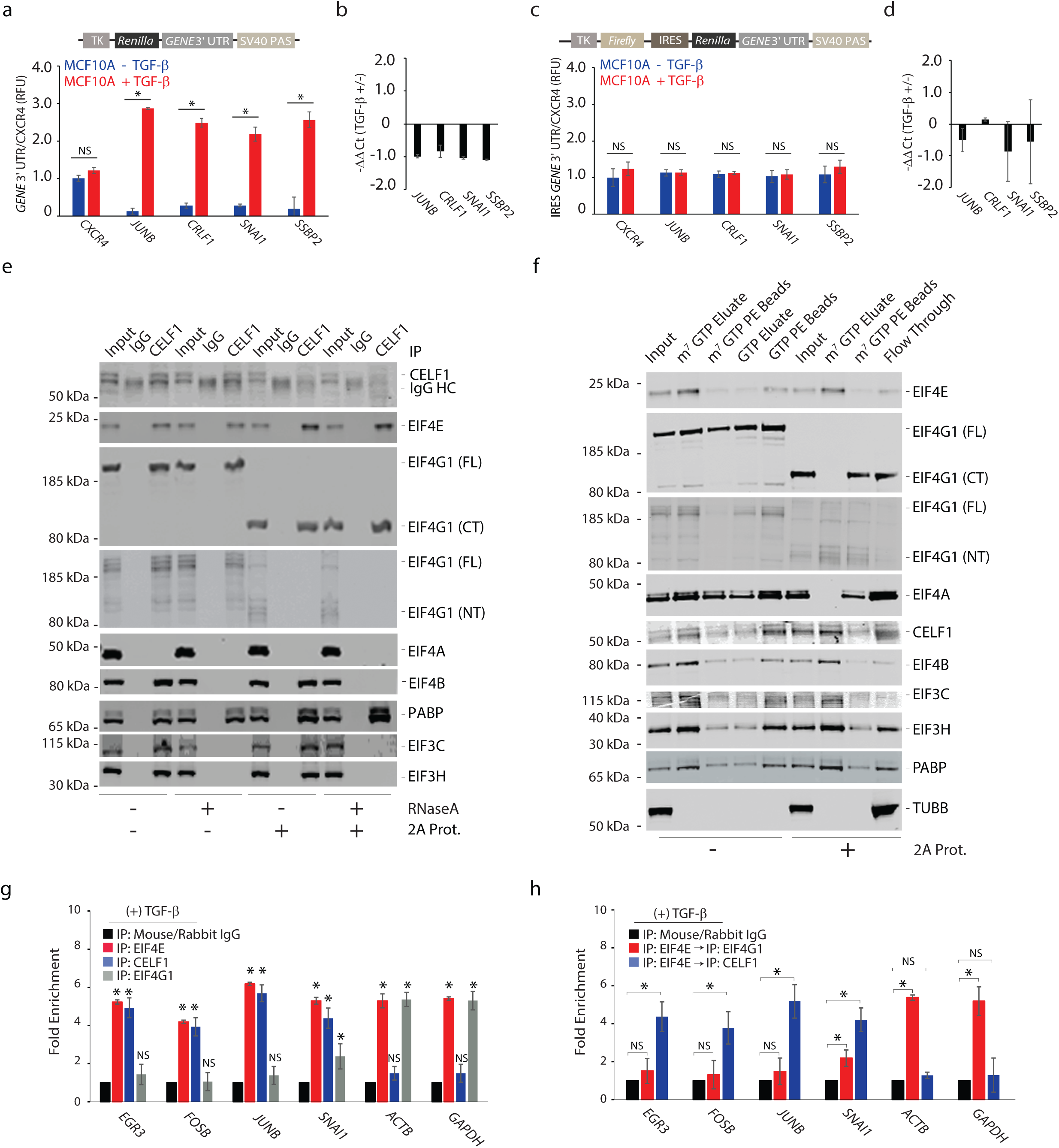
CELF interacts with eIF4E at the m^7^G cap, independent of intact eIF4G1. (**a**) Reporter assay quantifying the relative Renilla luciferase expression from the indicated 3’ UTR luciferase reporters in untreated and TGF-β-treated MCF10A cells. Data was normalized to Firefly luciferase expression and presented as fold change relative to CXCR4 reporter in untreated MCF10A cells. (**b**) Quantitative real-time PCR (qRT-PCR) of indicated *Renilla* luciferase reporters (pRL-TK) in untreated and TGF-β-treated MCF10A cells. Data was normalized to *ACTB* expression. (**c**) As ***a***, with the indicated 3’ UTR luciferase reporters driven from an internal ribosomal entry site (IRES) in untreated and TGF-β treated MCF10A cells. (**d**) As ***b***, for reporter assays in ***c***. (**e**) Left six lanes - Immunoblots of immunoprecipitates from lysates derived from MCF10A cells treated with TGF-β for 72 hours. One half of each total immunoprecipitate was digested with RNase A prior to immunoblotting with the indicated antibodies. Right six lanes – As in the left six lanes, but lysates were digested with coxsackievirus 2A protease to cleave eIF4G1 before immunoprecipitation. CT =C-terminal; FL =full length; NT = N-terminal. (**f**) m^7^GTP cap analog binding assays utilizing cytosolic extracts derived from MCF10A cells treated with TGF-β for 72 hours. As above, one half of each extract was digested with coxsackievirus 2A protease to cleave eIF4G1 before the assay. (**g**) RNA crosslinking-immunoprecipitation/qRT-PCR of GRE-containing mRNAs (*EGR3, FOSB*, *JUNB*, *SNAI1*) from TGF-β treated MCF10A cells using anti-CELF1, anti-eIF4E, anti-eIF4G1 antibodies or mouse and rabbit IgG. *ACTB* and *GAPDH* are non-GRE containing negative control mRNAs. (**h**) RNA crosslinking-immunoprecipitation/qRT-PCR of GRE-containing mRNAs (*EGR3, FOSB*, *JUNB*, *SNAI1*) from TGF-β treated MCF10A cells using tandem anti-eIF4E/anti-CELF1, anti-eIF4E/anti-eIF4G1 antibodies or mouse and rabbit IgG. *ACTB* and *GAPDH* are non-GRE containing negative control. In all panels, results are representative of at least three independent experiments and error bars depict mean ± standard deviation (SD) of aggregate replicates performed in triplicate. NS: not significant; *pval ≤ 0.05 (Student’s t-test).

We next assessed whether CELF1 physically interacts with established components of the canonical translation initiation complex. EMT was induced in MCF10A cells via either TGF-β treatment or *GFP-CELF1* overexpression and monitored by loss of expression of the epithelial cell marker E-cadherin (CDH1) concomitant with induction of expression of the mesenchymal cell markers Fibronectin (FN1) and Vimentin (VIM) (**Supplementary Figure 1a**). Except for induction of phosphorylation of eIF4E at serine 209, which has been previously shown as required during TGF-β-induced EMT (20), no change in steady state protein expression of any of the tested translation initiation factors was observed between the epithelial and mesenchymal states (**Supplementary Figure 1a**). Co-immunoprecipitation experiments using extracts from TGF-β-induced cells revealed an association among CELF1, eIF4E, eIF4G1, eIF4B, eIF3H, eIF3C, and PABP (Figure 1e). CELF1 was not physically associated with eIF4A, eIF4G2, eIF4G3, or eIF4E-binding protein (4E-BP) in any of the conditions assayed (Figure 1e and **Supplementary Figure 1b**). CELF1 retained its physical association with eIF4E, eIF4G1, and PABP following RNase A digestion, whereas this digestion eliminated the association of CELF1 with eIF4B, eIF3H, and eIF3C, consistent with the notion that these latter interactions are RNA-dependent (Figure 1e).

Given the established physical association of eIF4E, eIF4G1, and PABP the above results are consistent with a model in which CELF1 directly binds one or more of these three proteins. To differentiate among the inherent possibilities, we again immunoprecipitated cellular extracts derived from TGF-β treated MCF10A cells that were either untreated or digested with purified coxsackievirus 2A protease. Coxsackievirus 2A protease impairs m^7^G cap-dependent translation during the coxsackievirus replication cycle by cleaving the eIF4E and poly(A)-binding protein (PABP)-binding N-terminal fragment of eIF4G1 from eIF4G1’s eIF4A- and eIF3-binding C-terminal fragment (21–24). Following digestion with 2A protease, CELF1’s interaction between CELF1 with the C-terminal, but not N-terminal, fragment of eIF4G1 was maintained. Surprisingly however, the association between CELF1 and both eIF4E and PABP were preserved within this context (Figure 1e), demonstrating that this tripartite interaction was independent of intact eIF4G1. EIF3C, eIF3H, and eIF4B retained their association with CELF1 upon 2A protease digestion, unless also digested with RNase A (Figure 1e), indicating that these factors are a stable part of a remaining RNA-dependent complex that is not dependent on intact eIF4G1. We concluded that CELF1 is likely to bind eIF4E and PABP directly and that these associated proteins may be tethered to eIF4B and eIF3 in an RNA-dependent but eIF4G1-independent fashion.

While our experiments assessed CELF1’s putative interactions with members of the canonical translation pre-initiation complex, these interactions were in the context of whole cell lysate. We thus next sought to determine whether these interactions were preserved at the mRNA m^7^G cap, using direct capture of eIF4E and its binding partners using extracts from TGF-β-induced cells with cap-analog affinity resin and immunoblot analysis. Competitive elution with m^7^GTP, but not GTP, confirmed an association between eIF4E and CELF1, eIF4G1, eIF4A, eIF4B, eIF3H, eIF3C, and PABP at the m^7^G cap (Figure 1f). Following digestion with 2A protease, the N-terminal, but not C-terminal, fragment of eIF4G1 precipitated with m^7^G cap-analogue, co-eluting with eIF4E as expected (22–24) (Figure 1f). CELF1’s association with the m^7^G cap-analogue was sensitive to competition with m^7^GTP but not GTP, and this association was preserved in both the presence and absence of intact eIF4G1 (Figure 1f).Digestion with 2A protease also disrupted eIF4A’s interaction with the m^7^G cap, but did not inhibit interaction of eIF4B, eIF3C, eIF3H, and PABP (Figure 1f). However, the interaction between eIF4G1’s C-terminus and CELF1 observed in whole cell lysate (Figure 1e) was not recapitulated in the context of cap-analogue binding (Figure 1f). This suggested that the CELF1/eIF4E/PABP interaction at the m^7^G cap of mRNA is likely to be independent of a direct association between CELF1 and eIF4G1.

To further test this model, we performed UV crosslinking/immunoprecipitation/qRT-PCR (RIP) assays from TGF-β-treated cells (Figure 1g). Immunoprecipitation from cellular extracts derived from these cells with anti-CELF1 antibody revealed significant enrichment of GRE-containing (*EGR3*, *FOSB*, *JUNB*, and *SNAI1*) but not control (*GAPDH* and *ACTB*) mRNAs. As expected, immunoprecipitation with anti-eIF4E antibody resulted in significant enrichment for each of the mRNAs tested. In contrast, while immunoprecipitation with anti-eIF4G1 enriched each of the control mRNAs, this immunoprecipitation enriched only one GRE-containing mRNA (*SNAI1*). Similar results were obtained using tandem co-immunoprecipitation RIP assays in which anti-eIF4E immunoprecipitates were then immunoprecipitated once more with anti-CELF1 or anti-eIF4G1 (Figure 1h).

### CELF1 reduces the necessity for eIF4G1 in translation of GRE-containing EMT effector mRNAs

To directly assess the relationship between whether CELF1 and eIF4G in promoting the translation of GRE-containing mRNAs, we performed *in vitro* luciferase assays using translationally competent extracts derived from TGF-β-treated MCF10A cells that had been transiently transfected with control shRNA targeting β-galactosidase (*GLB1* – negative control) or *CELF1*. As expected, robust translation of a capped and polyadenylated *Renilla* luciferase reporter mRNA generated from the *pRL-TK-CXCR4-6x* reporter plasmid (*CXCR4*) within these extracts was significantly attenuated following cleavage of eIF4G1 by 2A protease (p<0.01; Figure 2a; **Supplementary Figure 2a**). A *Renilla* luciferase reporter mRNA driven by the EMCV, but not the Hepatitis C virus (HCV) IRES, was impacted by 2A protease digestion (Figure 2a), consistent with published results (25–27). In contrast, cleavage of eIF4G1 had no significant effect on the translation of reporter mRNAs corresponding to established CELF1 target 3’ UTRs (*CRLF1* and *SNAI1*) if the native GRE was present within these UTRs (p<0.01; Figure 2a; **Supplementary Figure 2a**). Deletion of the GRE rendered the reporters sensitive to 2A cleavage. Interestingly, performing this assay in extracts from cells in which *CELF1* had been knocked down by shRNA revealed a decrease in translation of the *CRLF1* and *SNAI1* reporters irrespective of 2A protease digestion (again, contingent upon the presence of the GRE within these reporters) that was restored by addition of recombinant CELF1 protein (p<0.01 compared to CELF1 depleted extracts; Figure 2b). This strongly suggests that the GRE within these reporters is repressive within the epithelial state unless directly bound by CELF1, and that binding of CELF1 to these elements promotes translation of GRE-containing mRNAs with a reduced requirement for intact eIF4G1. These results do not, however, rule out that the N- and C-terminal cleavage products of eIF4G1 may retain some function in promoting the translation of the GRE-containing reporters.

**Figure 2.**
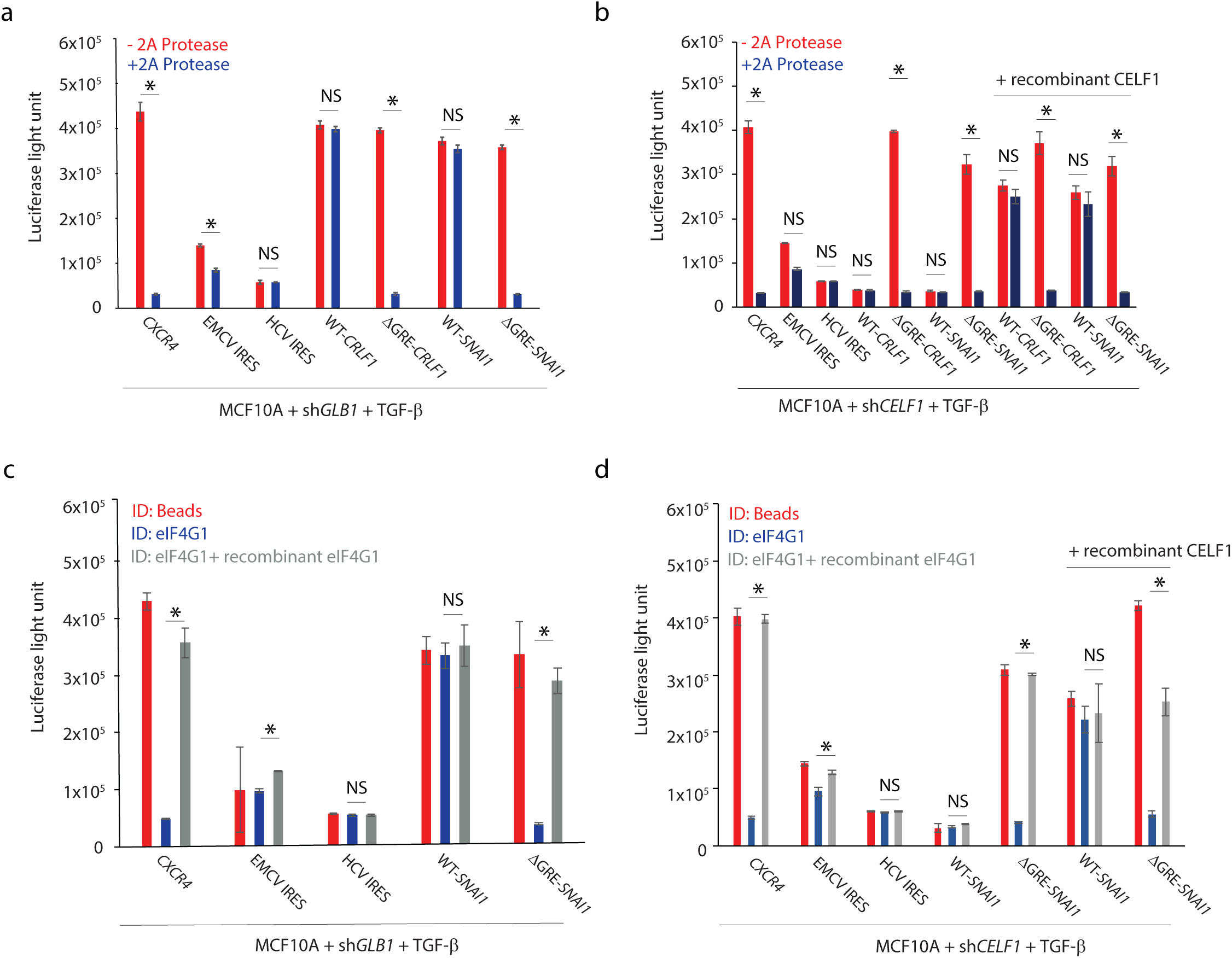
CELF1 stimulates translation of GRE-containing EMT effector mRNAs in the context of reduced eIF4G1 function. (**a, b**) Efficiency of *in vitro* translation of indicated capped and polyadenylated *Renilla* luciferase reporter mRNAs in mock or 2A protease-digested cell free extract. (**c, d**) Efficiency of *in vitro* translation of reporter mRNAs as described in (***a***) and (***b***), but with mock depleted (*beads*), eIF4G1 immunodepleted (*ID*) cell-free extract, or eIF4G1 immunodepleted extract reconstituted by addition of enriched eIF4G1. In (***a-d***), all extracts were derived from TGF-β-treated MCF10A cells transiently transfected with shRNAs targeting either *GLB1* (***a, c***) or *CELF1* (***b, d***). CXCR4 = cap-dependent control, EMCV IRES = eIF4G1-dependent but cap-independent control, HCV IRES = eIF4G1-and cap-independent control, WT = wild-type 3’ UTR, ΔGRE = 3’ UTR with deletion of GRE. In all panels, results are representative of at least three independent experiments and error bars depict mean ± standard deviation (SD) of aggregate replicates performed in triplicate. NS: not significant; *pval ≤ 0.05 (Student’s t-test).

To directly address this caveat, we repeated the *in vitro* translation assay using extracts from which eIF4G1 had been immunodepleted (**Supplementary Figure 2b**). As previously observed in the context of 2A protease digestion, immunodepletion of eIF4G1 had no effect on the translation of control reporters or reporters fused to GRE-containing CELF1 target UTRs (p>0.05), whereas translation of mutant UTRs lacking a GRE was significantly attenuated (p<0.01; Figure 2c and **Supplementary Figure 2e**). As expected, addition of recombinant eIF4G1 in excess to the depleted extracts restored translation of the control and Δ*GRE* reporters to levels matching what was observed in the mock-depleted extracts (Figure 2c and **Supplementary Figure 2c, e**). Again, *CELF1* knockdown diminished translation of the WT *SNAI1* and *CRLF1* reporters (p<0.01 in each case), even in extracts retaining eIF4G1 function, and this decrease in translation was rescued by addition of recombinant CELF1 Figure 2d and **Supplementary Figure 2d**, **f**). Our results are consistent with the notion that the GRE element inhibits translation of EMT effector mRNAs unless this element is bound by CELF1, which in turn reduces the dependency of translation of the GRE-containing mRNA on eIF4G1.

### Phosphorylation of eIF4E is required for CELF1-driven EMT in MCF10A cells

We next examined the direct interactions taking place within the CELF1-containing translation pre-initiation complex more closely. It has been previously established that phosphorylation of eIF4E is required for TGF-β-induced EMT in MCF10A cells (20), and CELF1 expression promotes the translation of EMT effectors at the level of translational initiation (Figure 1a-d). Given that our data suggests that CELF1 interacts with eIF4E at the m^7^G cap of GRE-containing mRNAs (Figure 1f), we asked whether disruption of eIF4E phosphorylation would be sufficient to block CELF1-driven EMT in MCF10A cells. We employed a well-established system (20) in which MCF10A cells are stably transduced with either wild-type or S209A phosphor-null mutant murine *Eif4e*, followed by shRNA-mediated knock down of endogenous human *EIF4E*. While cells expressing a wild-type *Eif4e* transgene underwent EMT normally in response to *CELF1* overexpression, *CELF1*-driven EMT was blocked by expression of the S209A mutant *Eif4e* (Figure 3a). Reciprocal co-immunoprecipitation experiments demonstrated that CELF1 interacts with eIF4E, and that this interaction is strengthened by eIF4E phosphorylation (Figure 3b). Indeed, translation of CELF1’s previously defined EMT effector targets (10) was blocked in the S209A mutant expressing cells (Figure 3c-e), consistent with the notion that eIF4E phosphorylation is required for CELF1’s function as a promoter of mRNA translation and EMT driver.

**Figure 3.**
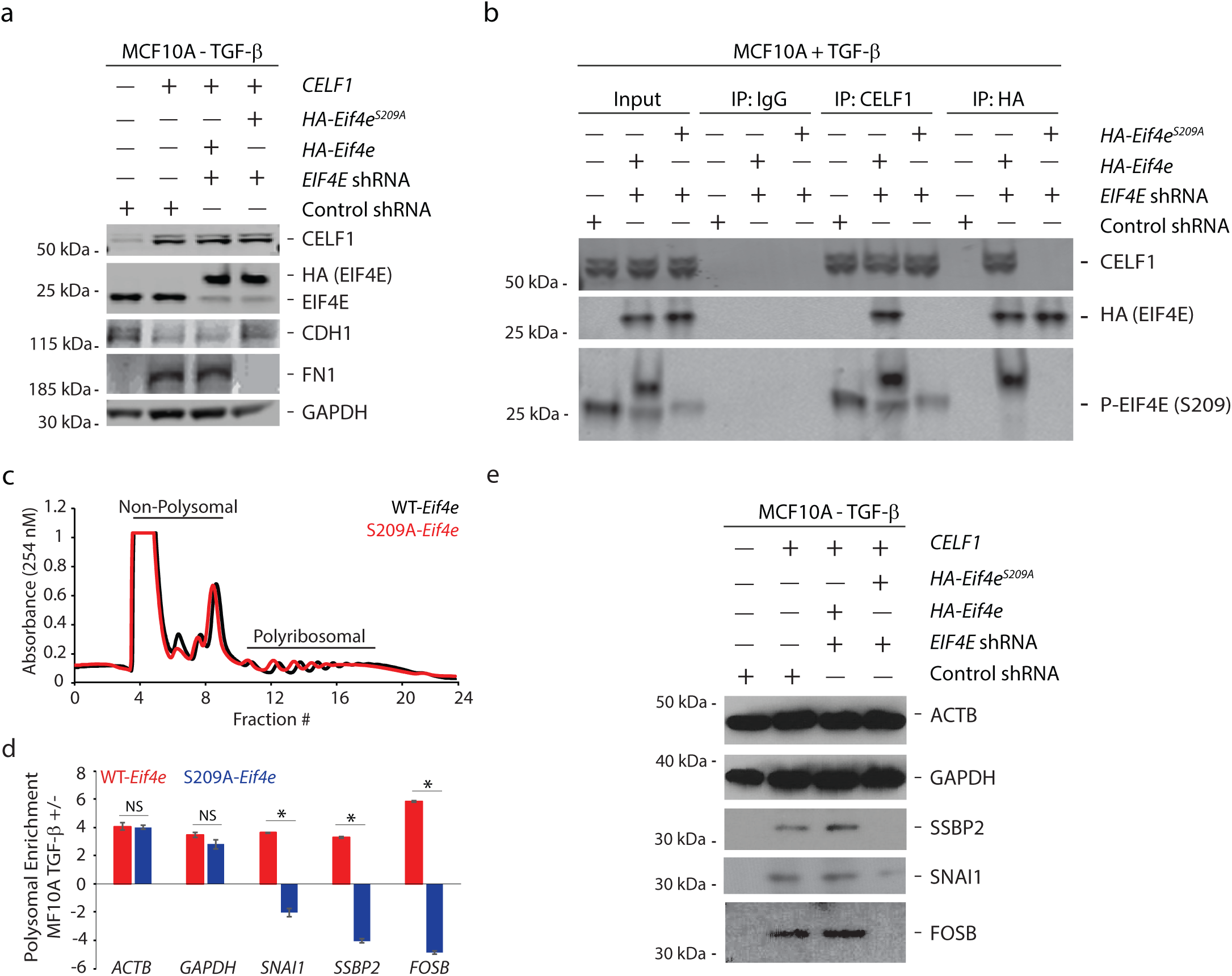
Phosphorylation of eIF4E is required for CELF1-driven EMT in MCF10A cells. (a) Phosphorylation of eIF4E is required for CELF1-driven EMT in MCF10A cells. **(a)** Immunoblots of lysates derived from MCF10A cells stably expressing either HA-tagged wild-type or S209A mutant murine *Eif4e* and shRNA targeting human *EIF4E* or control shRNA, and either mock transfected or transiently transfected with a CELF1 overexpression construct for 72 hours. GAPDH = loading control. **(b)** Immunoblot of immunoprecipitates from lysates derived from TGF-β-treated MCF10A cells, stably expressing either HA-tagged wild-type or S209A mutant murine *Eif4e* and shRNA targeting human *EIF4E* or control shRNA. IgG: negative immunoprecipitation control. **(c)** Polysomal profiles from MCF10A cells in which endogenous *EIF4E* expression had been knocked down via shRNA and then rescued via stable transduction of either wild-type (WT) or an S209A mutant *Eif4e*. **(d)** Quantitative real-time PCR (qRT-PCR) validation of polyribosomal enrichment and depletion of indicated mRNAs using total and polyribosomal mRNA in TGF-beta-treated MCF10A cells stably expressing wild-type (WT) or S209A mutant *Eif4e*. **(e)** MCF10A cells in which endogenous *EIF4E* expression had been knocked down via shRNA and then rescued via stable transduction of either wild-type (WT) or an S209A mutant *Eif4e* were transiently transfected with CELF1 expression construct and assessed via immunoblot for CELF1-regulated EMT effectors after 72 hours. In all panels, results are representative of at least three independent experiments and error bars depict mean ± standard error of mean (SEM) of aggregate replicates performed in triplicate. NS: not significant; *pval ≤ 0.05 (Student’s t-test).

### CELF1 is an eIF4E-binding protein (4E-BP)

We next sought to further establish the direct interactions CELF1 might make within the translation pre-initiation complex. The YXXXXLφ motif (where X is any amino acid and φ is a hydrophobic amino acid residue) is common among eIF4G proteins and 4E-BPs, which utilize the motif to compete for binding to the tryptophan (W) 73 residue on the dorsal surface of eIF4E (28). We identified three putative eIF4E binding motifs in the CELF1 peptide sequence (Figure 4a), and transiently transfected MCF10A cells with GFP tagged wild-type *CELF1* or mutants of *CELF1* in which these putative eIF4E-binding motifs were deleted, either individually or in combination. Immunoprecipitation of WT CELF1 or these CELF1 mutants revealed that deletion of broadly conserved CELF1 amino acids 365-371 (YAAAALP; **Supplementary Figure 3a**) abrogated precipitation of eIF4E (Figure 4b). Similar results were obtained using affinity-tagged purified proteins (Figure 4c and **Supplementary Figure 3b**), establishing that CELF1 interacts directly with eIF4E and may compete with 4E-BPs and eIF4G1 for binding to W73 on the latter protein. Importantly, CELF1’s direct interaction with eIF4E was independent of previously described post-translational modifications (PTMs) (29) (**Supplementary Figure 3c-e**), even though mutations abrogating these features within CELF1 disrupted *CELF1* overexpression-driven EMT (**Supplementary Figure 3f**).

**Figure 4.**
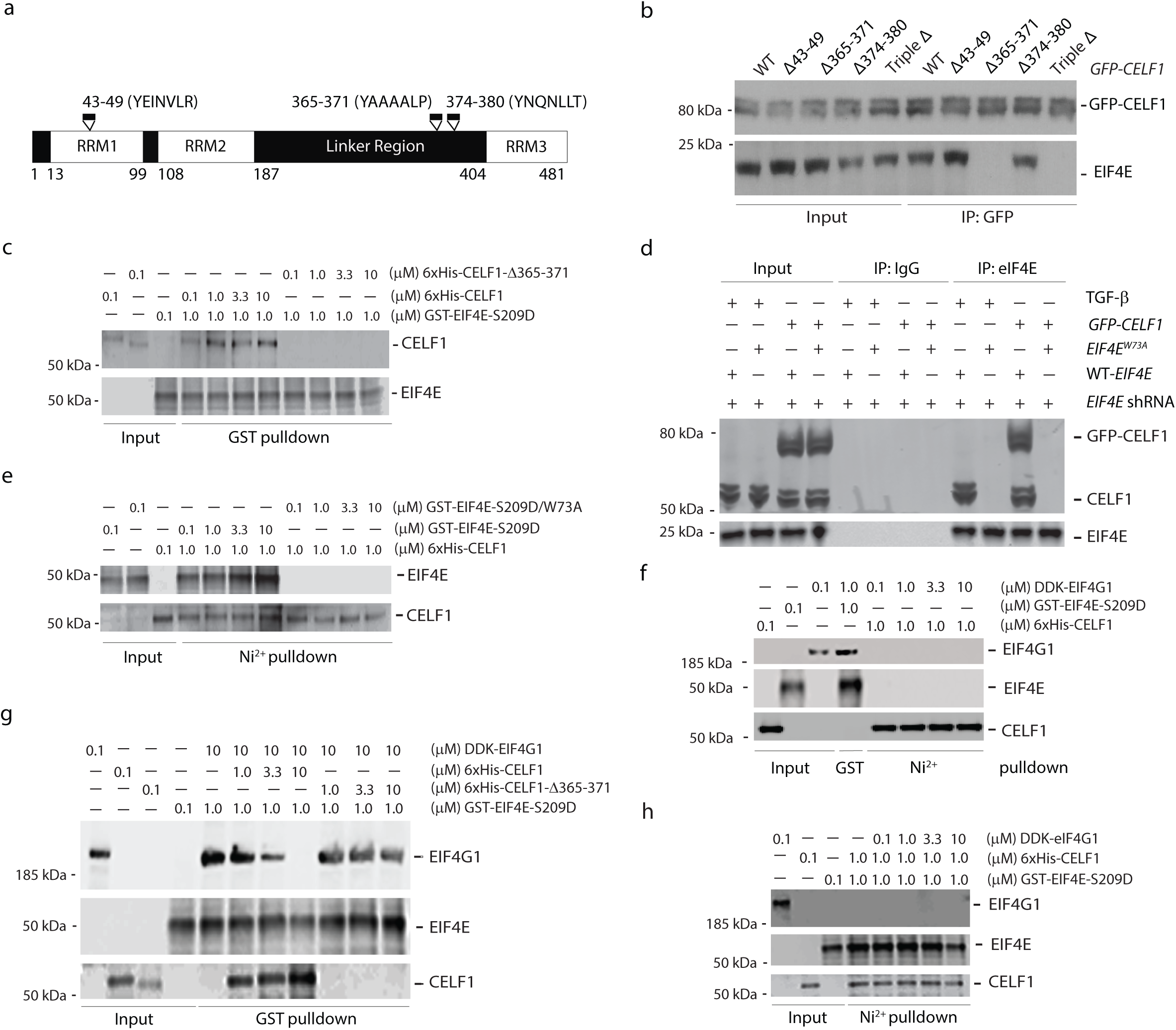
CELF1 is a 4E-BP and directly binds eIF4E at tryptophan 73. (**a**) Schematic of CELF1 domain structure and candidate eIF4E binding motifs. RRM, RNA-recognition motif. (**b**) Immunoblots of immunoprecipitations derived from lysates of MCF10A cells transfected with WT or indicated mutant *GFP-CELF1* plasmids for 72 hours. (**c**) Immunoblots of binding assays using affinity-purified phosphomimic eIF4E (*GST-eIF4E S209D*), affinity-purified wild type CELF1 (*6xHis-CELF1*) or affinity-purified mutant CELF1 (*6x-His-CELF1-Δ365-371*). (**d**) Immunoblots of immunoprecipitations derived from lysates of MCF10A cells stably expressing a shRNA targeting the 3’ UTR of *EIF4E* and co-expressing either wild-type or W73A mutant *EIF4E*, and transiently transfected with a *GFP-CELF1* plasmid for 72 hours. IgG: negative control. (**e**) Immunoblots of binding assays in which affinity-purified 6xHis-CELF1 was mixed with affinity-purified phosphomimic (*GST-eIF4E-S209D*) or mutant (*GST-S209D/W73A*) eIF4E. (**f**) Purified 6xHis-CELF1 or GST-eIF4E-S209D was mixed with purified DDK-eIF4G1. Complexes were pulled down with Ni^2+^ or purified GST beads, respectively, and immunoblotted with indicated antibodies. (**g**) Immunoblots of binding assays in which purified GST-eIF4E S209D was mixed with enriched DDK-eIF4G1 in combination with either purified wild type (*6xHis-CELF1*) or mutant CELF1 (*6x-His-CELF1-Δ365-371*). (**h**) Immunoblots of binding assays in which purified GST-eIF4E-S209D was mixed with either purified 6xHis-CELF1 or purified 6xHis-CELF1 in combination with purified DDK-eIF4G1. All results are representative of at least three individual experiments.

Given the necessity of the 4E-BP like consensus sequence for CELF1’s binding to eIF4E, we next asked whether this interaction was analogously mediated by the same W73 residue bound by eIF4G1 and other 4E-BPs. We transduced MCF10A cells with either wild-type or W73A mutant *EIF4E*, followed by knock down of the endogenous *EIF4E* using shRNA targeting the 3’UTR of *EIF4E*. Similar to previous descriptions (30–32), stable overexpression of the W73A mutant *EIF4E* did not markedly inhibit global translation in this context as assessed by the surface sensing of translation using puromycin labeling (SUnSET) assay (**Supplementary Figure 3g**) (33). Immunoprecipitation of WT eIF4E or the W73A mutant eIF4E from MCF10A extracts revealed that CELF1’s interaction with eIF4E was dependent upon the W73 residue (Figure 4d). Our strategy of expressing the W73A mutant *EIF4E* in MCF10A cells precluded us from directly testing successful knockdown of the endogenous *EIF4E*. That we did not observe any interaction of endogenous CELF1 or ectopically overexpressed GFP-CELF1 with eIF4E in MCF10A cells expressing W73A mutant *EIF4E* confirmed successful knockdown of endogenous *EIF4E* in these cells. Again, these results were recapitulated using affinity-tagged purified proteins, indicating that the W73 mutation ablated the direct interaction between CELF1 and eIF4E in this latter context (Figure 4e). Taken together with the above results, these data suggest that CELF1 is a novel mammalian 4E-BP that promotes translation of GRE-containing mRNAs in mesenchymal cells.

As stated above, given that canonical 4E-BPs block translation by competing with eIF4G1 for binding to eIF4E, it was counterintuitive that CELF1 would use a similar mechanism to promote translation. We thus sought to further investigate associations among the binding of eIF4E, CELF1, and eIF4G1 using affinity-purified proteins. Consistent with established literature and our own data described above, *in vitro* binding assays confirmed that eIF4G1 binds wild-type eIF4E, but neither the eIF4E W73A mutant (**Supplementary Figure 4a**) nor CELF1 (Figure 4f). In addition, while purified CELF1 was able to stoichiometrically compete with purified eIF4G1 for eIF4E binding (and this competition was dependent upon CELF1 amino acids 365-371 - Figure 4g), eIF4G1 was not conversely able to displace wild-type CELF1 protein bound to eIF4E (Figure 4h). Additional *in vitro* binding experiments confirmed that CELF1 does not directly bind eIF4A (**Supplementary Figure 4b**).

We next explored interactions among eIF4G1, CELF1, and PABP, again using affinity-purified proteins. As expected, *in vitro* binding confirmed interaction of PABP and eIF4G1, and consistent with our experiments performed using cell extracts, we found that CELF1 can also directly PABP (**Supplementary Figure 5c**). Interestingly, competitive binding assays revealed that eIF4G1 did not displace CELF1 from PABP, but was instead able to bind the two complexed proteins (**Supplementary Figure 4c**). These results indicate that CELF1 directly binds PABP and can thus in theory promote mRNA circularization of GRE-containing targets, but that it does this by binding a site on PABP distinct from the site bound by eIF4G1.

### Interaction of CELF1 and eIF4E is required for CELF1-driven EMT in vitro and experimental metastasis in vivo

Given that we had previously demonstrated that overexpression of CELF1 is sufficient to induce EMT in several breast epithelial cell lines, we next set out to determine whether disruption of CELF1’s interaction with eIF4E would abrogate this induction. Indeed, overexpression of the *CELF1*Δ365-371 (YAAAALP) mutant failed to induce EMT in MCF10A cells despite comparable levels of expression between the mutant and wild-type CELF1 proteins (Figure 5a). Conversely, a W73A mutation within the eIF4E sequence disrupted CELF1’s ability to induce EMT in MCF10A cells (Figure 5b). To further generalize this observation, we examined the isogenic derivative of the parental non-transformed MCF10A line, the non-malignant derivative line, MCF10AT1. We previously demonstrated that treatment of the MCF10AT1 cells with TGF-β for 72 hours resulted in the increased expression of CELF1 protein and EMT, and overexpression of *CELF1* promoted EMT in MCF10AT1 cells independently of TGF-β (10). Hence, we tested if overexpression of *CELF1* Δ365-371 (YAAAALP) mutant would fail to induce EMT in the MCF10AT1 cells. EMT of the transductants was monitored via immunoblot analysis of molecular markers and migration and invasion in standard transwell assays. Whereas overexpression of wild-type *GFP-CELF1* induced the mesenchymal cell markers FN1 and VIM and loss of epithelial cell marker CDH1, overexpression of the *CELF1* Δ365-371 (YAAAALP) mutant failed to induce EMT in the MCF10AT1 cells, as determined by the analysis of molecular markers (Figure 5c) and *in vitro* migration and invasion assays (Figure 5d-e). Similar results were obtained in the triple negative breast cancer cell line MDA-MB-468 (Figure 5c, f, g), indicating that interaction of CELF1 with eIF4E may be broadly required for EMT of breast epithelial cells.

**Figure 5.**
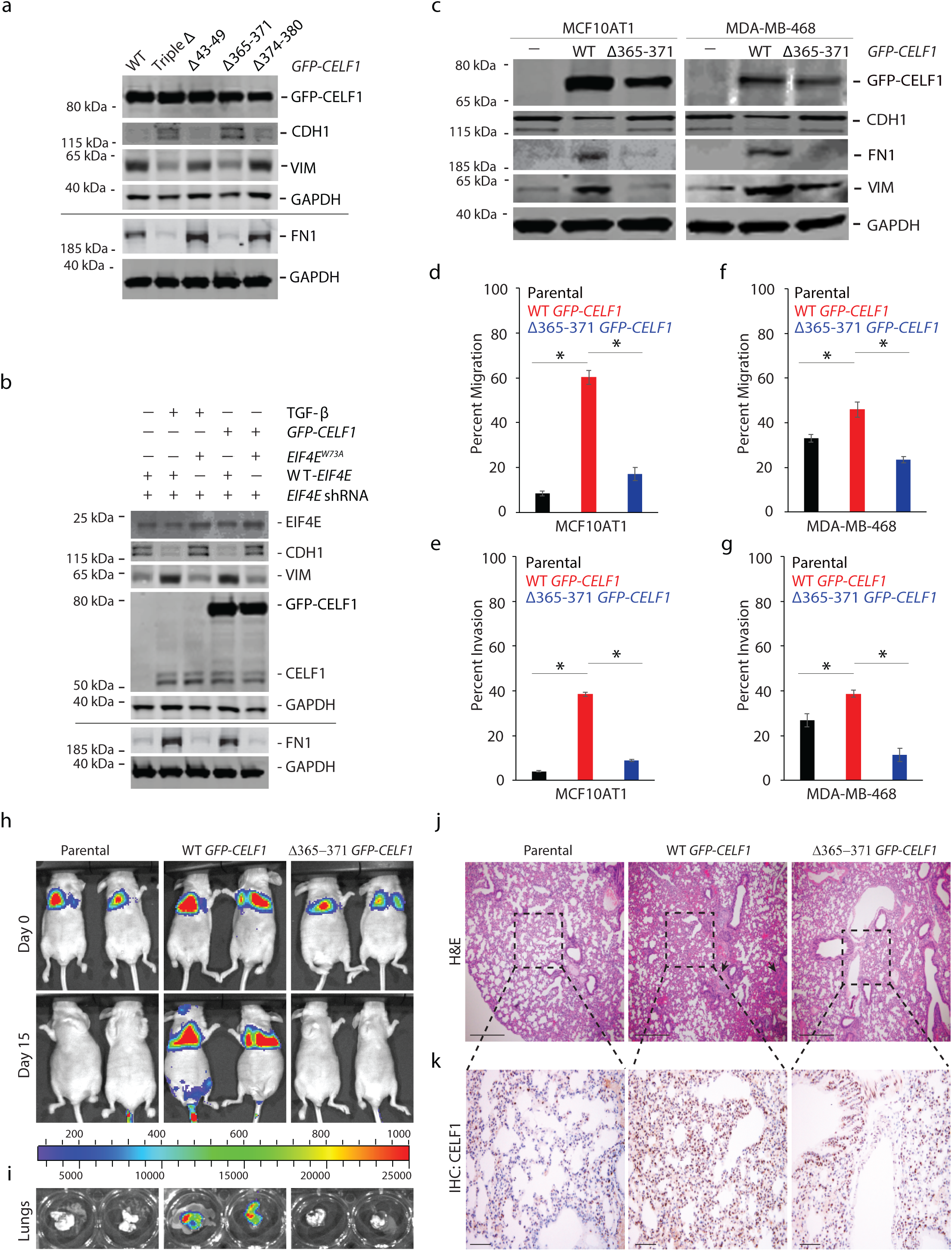
Interaction of CELF1 and eIF4E is required for CELF1-driven EMT and experimental metastasis. (**a**) Immunoblot analysis of indicated EMT markers in lysates derived from MCF10A cells transfected with WT or indicated mutant *GFP-CELF1* plasmids for 72 hours. (**b**) Immunoblot analysis of indicated EMT markers in lysates derived from MCF10A cells expressing either wild-type or W73A mutant human *EIF4E* and shRNA targeting the 3’ UTR of human *EIF4E* and induced to undergo EMT via stable expression of *GFP-CELF1* or TGF-β treatment for 72 hours. (**c**) Immunoblot analysis of indicated EMT markers and GFP-CELF1 in lysates derived from parental MCF10AT1 cells (*left column*) and MDA-MB-468 (*right column*), or each cell line stably transduced with either wild-type or Δ365-371 mutant *GFP-CELF1*. GAPDH = loading control in (***a***), (***b***) and (***c***); black line in (***a***) and (***b***) denote lysates derived from the same experiment, but gels processed in parallel. All results are representative of at least three individual experiments. (**d-g**) Quantification of relative *in vitro* cellular migration (***d***, ***f***) and invasion (***e***, ***g***) in transwell assays in parental MCF10AT1 and MDA-MB-468 cells, respectively, or stably transduced with either wild-type or Δ365-371 mutant *GFP-CELF1*. Data represents mean ± SD of at least three independent experiments, each performed in triplicates. *pval ≤ 0.05 (t-test). (**h**, **i**) Parental MCF10AT1 cells or stably overexpressing either wild-type or Δ365-371 mutant *GFP-CELF1* were injected into the tail vein of athymic nude mice. The incidence and progression of metastasis was measured by luciferin injection and bioluminescence imaging of Firefly luciferase (**h**), and *ex vivo* excised lungs on day 15 (**i**). (**j**, **k**) Representative hematoxylin and eosin (H&E) (**j**) and immunohistochemical (IHC) (**k**) staining, respectively, of the lungs from mice shown in (**h**). Scale bar, 200 µm (**j**); 50 µm (**k**). Black arrows (**j**) indicate micrometastases. Dotted lines indicate area shown in corresponding H&E staining of serial sections shown in (**j**). For (**h**-**k**), representative images are from n=4 for parental, n=4 for wild-type *GFP-CELF1*, and n=6 for Δ365-371 mutant *GFP-CELF1* experimental groups, respectively.

To determine whether the importance of the CELF1/eIF4E interaction was limited to EMT *in vitro*, we next assessed the importance of this interaction in a model of experimental metastasis. Overexpression of *CELF1* induces experimental lung metastasis from the normally tumorigenic but non-metastatic MCF10AT1 cell line (10,34). Consistent with our own previously described work, ectopic expression of wild-type *CELF1* within MCF10AT1 cells injected into the tail vein of nude mice conferred these cells with the ability to rapidly induce lung colonization in the animals tested. However, lung colonization was attenuated within MCF10AT1 cells overexpressing the *CELF1* Δ365-371 (YAAAALP) mutant (Figure 5h-k). In aggregate, our results strongly support a model in which within the context of breast epithelial cells undergoing EMT and metastatic dissemination, CELF1 is recruited to GRE-containing EMT effector mRNAs, acting in *cis* on these mRNAs to promote non-canonical translation of these mRNAs via a direct interaction with eIF4E and PABP (Figure 6).

**Figure 6.**
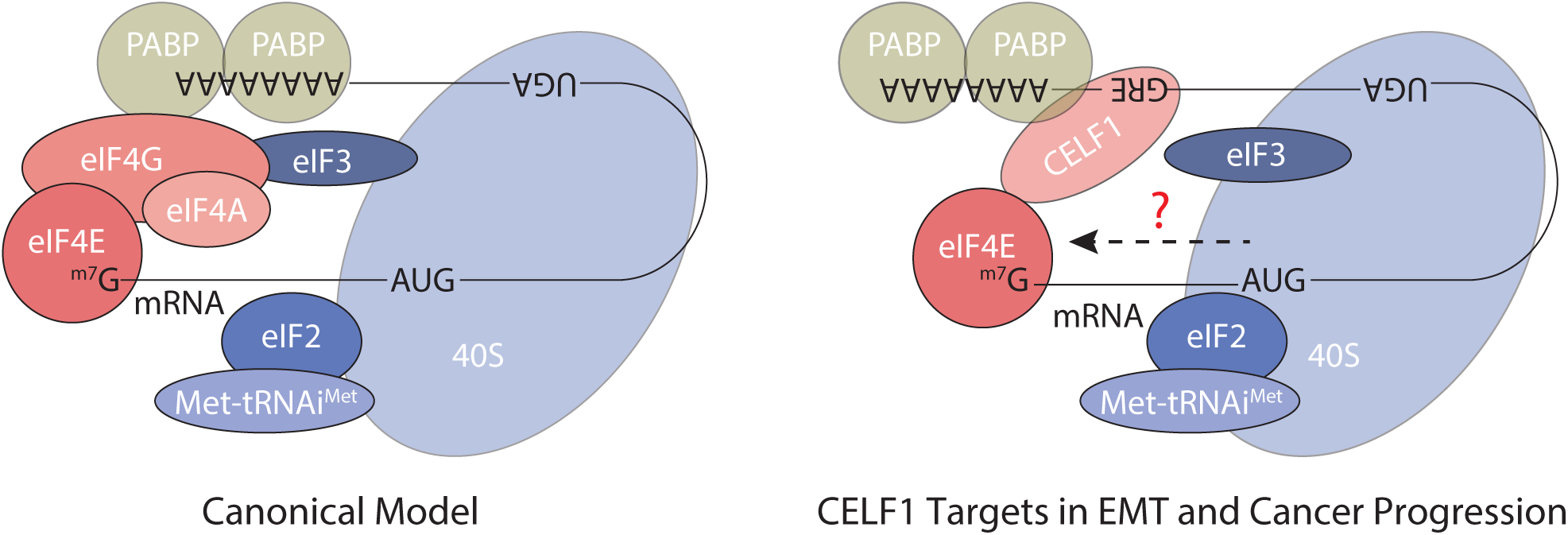
Working model of CELF1-dependent cap-dependent translational initiation during TGF-β-mediated EMT of breast epithelial cells. During EMT of breast epithelial cells, CELF1 is stabilized and recruited to GREs within the 3’ UTRs of EMT effector mRNAs. In *cis* on the RNA, CELF1 then bridges eIF4E and PABP to promote translation of these mRNAs. Additional mechanisms conferring specificity on this process as well as the mechanism by which CELF1 recruits the full 43S pre-initiation complex remain to be defined. Components are not drawn to scale, and eIF2 and eIF3 are multiprotein complexes depicted as single entities for simplicity.

## Discussion

Our results identify a cap-dependent non-canonical translation initiation mechanism that facilitates EMT and metastatic progression by selectively promoting translation of GRE-containing EMT effector mRNAs. Even though the eIF4F complex has long been considered as core to canonical cap-dependent translation initiation (35), there is a developing paradigm that cap-dependent translation initiation independent of eIF4F may exist as a cellular mechanism for adaptive translation. This is especially true in the context of physiological or pathological conditions where canonical cap-dependent translation is broadly inhibited (36–40). However, to our knowledge a mechanism in which a 4E-BP directly associates with a *cis*-element in a mRNA’s 3’ UTR and is thus incorporated into a specialized complex to promote (rather than inhibit) the translation of the associated mRNA, has not been previously described. CELF1 itself has been previously implicated in regulation of the translation of the C/EBPβ transcription factor (41) and p21 protein (42) in hepatocytes and fibroblasts, respectively. However, the mechanism described for this regulation in these contexts is markedly different from the mechanism we have defined here. In both of these previous reports, CELF1 mediates regulation of translation by binding near the 5’ region of the C/EBPβ and p21 coding sequences, and an interaction between CELF1 and the eIF2 initiation factor has been implicated in CELF1’s regulation of C/EBPβ (43). In contrast, our data suggest that CELF1-mediated regulation of its targets in the context of EMT is via recruitment to GREs in the 3’ UTR of affected transcripts, and we do not observe co-immunoprecipitation of CELF1 and eIF2 within our own experimental model system (data not shown).

In the scanning model of translation initiation, the decoding site and latch of the 40S subunit must open to allow the recruitment and migration of mRNA, and this unwinding process is catalyzed by the eIF4A helicase (11). Surprisingly, our results support the conclusion that CELF1’s role in promoting translation of GRE-containing EMT effector mRNAs is independent of an interaction with eIF4A. Much work has led to the now generally accepted notion that the 5’ UTRs of mRNAs encoding oncogenic factors are characterized by increased length and/or more complex structure, and thus that translation of these mRNAs is more dependent upon eIF4A helicase activity than translation of mRNAs whose 5’ UTRs are less structured (44–46). Importantly, most of the studies establishing this notion have focused specifically on mRNAs encoding regulators of the cell cycle rather than EMT or de-differentiation (38–40,47). This, taken together with the observations described within several recent published studies highlighting non-canonical modes of translation initiation in transformed cells (39,40,47), engender speculation that different facets of tumorigenesis or tumor progression incorporate discrete modes of canonical or non-canonical translation initiation specific to these facets. More broadly however, it has also been shown that chemical inhibition of eIF4A does not significantly affect *in vitro* survival of breast epithelial cells - and indeed upregulates translation of certain transcripts (48) - supporting the notion that robust translation initiation mechanisms that are not dependent upon eIF4A activity exist within these cells. Interestingly, recent work in yeast has implied that eIF4A’s cofactor eIF4B may function as an eIF4A-independent helicase for mRNAs with long structured 5’ UTRs but weak dependence on eIF4G (49). Our demonstration here that CELF1 interacts with eIF4B may hint that a similar mechanism may be at play within our model system, but formal testing of this hypothesis, or identification of the helicase functioning within this context, is beyond the scope of the current study.

Perhaps even more surprising than the apparent absence of eIF4A in CELF1-dependent translation complexes is the finding that CELF1 can be classified as a 4E-BP that stimulates translation initiation. Virtually all 4E-BPs described to date bind eIF4E at W73 residue, thereby sterically occluding the eIF4E/eIF4G interaction, formation of the eIF4F translation pre-initiation complex, and bulk translation. Given this broadly conserved mechanism, our finding that CELF1, brought in *cis* by GRE-containing mRNAs, by definition occludes eIF4G by binding W73 yet concomitantly promotes translation of these mRNAs, is to our knowledge without precedent within a mammalian system. To our knowledge, the only other example of a 4E-BP able to coordinate translation initiation is *Mextli*, a gene product identified in *Drosophila* that promotes germ stem cell maintenance (50). Nonetheless, it is intuitively clear that additional regulatory mechanisms are likely to dictate CELF1’s recruitment to the translation pre-initiation complex and its activity within that complex, and that unraveling these mechanisms is an exciting arena for future study.

The bulk of our functional data might be interpreted to suggest that CELF1-mediated translation of GRE-containing EMT effector mRNAs is independent of eIF4G1. However, it should be noted our methodology does not entirely rule out a role for eIF4G1 given that residual protein remains after immunodepletion and perhaps even after cleavage with 2A protease. Indeed, the global function of the core translation machinery has been consistently shown to be surprisingly refractive to depletion of central components of this machinery (32,36–40,51). Additionally, our data does not formally rule out that eIF4G1 is incorporated into CELF1-conatining translation initiation complexes in a non-canonical fashion. Significant additional work is required to fully characterize the components of the CELF1-specific translation initiation complex within this context and determine precisely how the 43S pre-initiation complex is recruited to these mRNAs. In addition, given that our results demonstrate an RNA-independent interaction with eIF4E and PABP, both of the latter of which are thought to bind virtually all mRNAs, it is patently clear that additional mechanisms dictating the specific incorporation of CELF1 into translation pre-initiation complexes of GRE-containing mRNAs remain to be discovered. Further, that GREs within the mRNAs that we have examined here confer repression in the epithelial state, where CELF1 protein is not present, suggests the existence of a second *trans*-regulator that must be evicted by CELF1 to allow translation activation and then EMT to occur. This is an exciting arena for future study.

While the components, mechanisms, and underlying signaling pathways dictating canonical translation pre-initiation complex formation in mRNA translation are well-established, the contribution of non-canonical complexes to translation is only beginning to be revealed. Already however, it is clear that regulation of translation is both context- and cell-type dependent in terms of target specificity and activity (39,40,47,52). In theory, encoding more than one mechanism of cap-dependent translation initiation allows cells additional control over protein synthesis, perhaps where additional granularity in this control might provide the cell an adaptive advantage in response to a specific physiological stimulus or genetic insult. It is thus tempting to speculate that additional specialized translational complexes remain to be discovered, both in normal cellular function and within the diseased or transformed state. From the latter perspective, a more robust mechanistic understanding of analogous specialized mechanisms of translation may reveal novel vulnerabilities that may be exploited in the clinic, especially given the initial promise within clinical trials of drugs targeting canonical translation initiation as a bulk process (53,54),

## Materials and Methods

### Cell culture and treatment

The MCF10A cell line was obtained from the ATCC (Manassas, VA) and cultured as described previously (10). MCF10A stable cells expressing HA-*eif4E* or HA-*eif4E^S209A^* were kind gifts from Dr. Wilson H. Miller, Jr. (Lady Davis Institute for Medical Research, Segal Cancer Centre, Jewish General Hospital; and McGill University; Montreal, Quebec, Canada) (20). MCF10A cells were treated for 3 days with 5 ng/ml of TGF-β1 (R&D Systems, Minneapolis, MN).

### Cell lysis, immunoblot, and immunoprecipitation

Cell lysis and immunoblotting was performed as described previously (10,55). **Supplementary Table 1** provides the list and associated information of antibodies used in the current study. All blots were also probed with GAPDH to confirm equal loading. For immunoprecipitation, cells were lysed with NP-40 lysis buffer (150 mM NaCl, 1% NP-40, and 50 mM Tris-Cl (pH 8.0) supplemented with protease and phosphatase inhibitor cocktail (ThermoFisher Scientific, Waltham, MA). Lysates (800 µg) were incubated at 4°C with 10 µg of indicated antibodies (**Supplementary Table 1**) or IgG (mouse or rabbit based on the antibody used for the immunoprecipitation) for 4 hours followed by addition of 20 μl protein A/G agarose beads (ThermoFisher Scientific) for 2 hours. Immune complexes were washed three times with constant rotation at 4°C (5 minutes each time) in lysis buffer. Beads were quickly centrifuged at maximum speed and immunoprecipitates were subjected to immunoblot analysis. Where indicated immunoprecipitates were treated with RNase A (100 µg per mg of protein lysate used for immunoprecipitation; Sigma-Aldrich, St. Louis, MO) (56), or recombinant coxsackievirus B3 2A protease (4 ng/µl) for 30 minutes at 37°C (57).

### RNA immunoprecipitation (RIP) and data analyses

RIP and data analyses was performed as described previously using primers described in Supplementary Table 2 (10).

### Polysome profiling, and quantitative real time PCR (qRT-PCR)

Polysome profiling from MCF10A cells stably expressing either wild-type or S209A mutant eIF4E and transiently transfected with *CELF1* expression plasmid (20) was performed using 10-50% sucrose gradients as described previously (10). Thirty OD units per condition were used for polysome profiling. We used TRIzol LS reagent (ThermoFisher Scientific) to extract RNA from equal volumes of the various polysome fraction and total lysate aliquots as described before (10). RNAs isolated from equal volume of polysome fractions or input total lysate were primed with random hexamers and reverse transcribed using SuperScript III Reverse Transcriptase (ThermoFisher Scientific). Control reactions omitted reverse transcriptase. The cDNAs were subsequently used for qRT-PCR reactions using KAPA SYBR FAST Universal 2X qPCR Master Mix (KAPA BIOSYSTEMS, Wilmington, MA) and the primers indicated in **Supplementary Table 2**. Data was plotted as the percent of total mRNA (% mRNA) in each fraction.

### m^7^GTP chromatography

Cytoplasmic extracts (10) were subjected to m^7^GTP chromatography using Immobilized γ- Aminophenyl-m^7^GTP (C10-spacer) beads (Jena Bioscience, Germany). Beads were equilibrated with buffer A (100 or 200 mM KCl, 50 mM Tris-HCl [pH 7.5], 5–10 mM MgCl_2_, and 0.5% Triton X-100) plus BSA (0.1 mg/ml) at 4°C for 30 minutes. The resin was washed and incubated with 500 µg of protein extract, and either left untreated or treated with 4 ng/µl of recombinant coxsackievirus B3 2A protease, for 60 minutes at 4°C. GTP (100 mM) was added to reduce nonspecific binding. The beads were washed with 0.4 ml of buffer A and then incubated for 30 minutes with 200 mM of m^7^GTP or GTP.

### Plasmid constructs

The pEGFP-N1-*CELF1* putative eIF4E binding mutants were generated via site-directed mutagenesis using the QuickChange II XL Site-Directed Mutagenesis Kit (Agilent Technologies, La Jolla, CA) and primers listed in **Supplementary Table 2**. For recombinant protein expression, the wild-type and Δ365-371 mutant *CELF1* were cloned in the pET His10 TEV LIC cloning vector (2B-T-10) (Addgene, Cambridge, MA) using NEBuilder HiFi DNA Assembly Master Mix (NEB) and primers listed in **Supplementary Table 2**. Other *CELF1* and *Renilla* luciferase expression plasmids utilized herein have been described before (10). The *GST-EIF4E* (wild-type and W73A mutant) cloned into the pGEX-5X-1 vector were kind gifts from Dr. Yan-Hwa Wu Lee (58) and were used to generate the phosphomimic wild-type and W73A mutant *GST-EIF4E* by site directed mutagenesis using primers listed in **Supplementary Table 2**. The wild-type and W73A *EIF4E* coding sequence entry constructs for overexpression were generated from FLAG-*EIF4E*, and FLAG-*EIF4E* (W73A) plasmids gifted by Dr. Katherine L Borden (59). Expression constructs for wild-type and W73A mutant *EIF4E* were generated by Gateway Cloning (ThermoFisher Scientific) into the *pLenti6.3* vector. ShRNA plasmid constructs targeting the 3’ UTR of *EIF4E* were cloned into the *pGIPZ* backbone using oligonucleotides listed in **Supplementary Table 2**.

### Transfection and transduction

Transient transfection was performed using Lipofectamine LTX (Life Technologies, Carlsbad, CA), per the manufacturer’s instructions. The *pGIPZ* lentiviral particles were generated by transfection of 293Ts using Lipofectamine 2000 (ThermoFisher Scientific), per the manufacturer’s instructions. The *pLenti6.3* lentiviral particles were generated by transfection of 293Ts as described previously(60). For transduction, early passage cells were seeded at 500,000 cells per 10-cm^2^ dish one day prior to infection and transduction was performed as described previously(61). To generate the *EIF4E* (W73A) overexpressing stable cells, the MCF10A cells were first transduced with the *pLenti6.3-EIF4E* (W73A) and selected with Blasticidin (5 µg/mL; R&D, Minneapolis, MN). Selected cells were then transduced with *pGIPZ* shRNA targeting the 3’ UTR of *EIF4E* and selected with Puromycin (2 µg/mL; R&D, Minneapolis, MN).

### Surface sensing of translation (SuNSET)

SuNSET technique was performed as described previously (33). Briefly, indicated cells were labeled for 10 minutes using puromycin (10 µg/ml). Cells were harvested, lysed as described above, and resolved using SDS-PAGE. Blots were probed with anti-Puromycin antibody (Kerafast, Boston, MA) and subsequently stained with Coomassie Blue R250 (Sigma-Aldrich).

### Recombinant protein expression and purification

The pGEX-5X-1 WT/S290D and W73A/S209D *EIF4E* constructs were transformed into competent BL21(DE3) *E. coli* (ThermoFisher). One colony was picked the following day, inoculated in 10 ml of Luria-Bertani (LB) broth and grown overnight (200 rpm at 37°C). The inoculate was added into 1000 ml LB (1:100) and grown at 200 rpm at 37°C for 2 to 3 hours until OD_600_ was between 0.6 and 0.8, at which point it was induced with 0.4 mM isopropyl β-D-1-thiogalactopyranoside (IPTG, ThermoFisher Scientific) and grown overnight at 200 rpm at 20°C. Following growth, cells were harvested and pelleted (5000 *g*, 10 minutes). The cell pellet was resuspended in 50 ml lysis buffer (50 mM PBS, pH7.6, 500 nM NaCl, 1 nM EDTA, and 1 mM PMSF) and disrupted by passing through microfluidizer. Lysate was clarified by centrifugation at 50,000 *g* for 30 minutes at 4°C. Clarified lysate was passed through a 022 µM syringe filter and loaded onto 1 ml GE GSTrap 4B column (GE AKTA) at 0.5 ml/minute. The column was washed with 20 column volumes of lysis buffer before elution using freshly prepared lysis buffer containing 20 mM glutathione at 0.5 ml/minute. The expression plasmids for the 6xHis-tagged CELF1 proteins were transformed, cultured, and induced similarly as above. The His SpinTrap Kit (GE Healthcare) was used according to manufacturer’s recommendations to purify the recombinant CELF1 proteins. Finally, buffer exchange was performed using 5 ml desalting spin-columns (MilliporeSigma, Burlington, MA). Desalted and purified protein samples were aliquoted, glycerol was added at 10% (v/v), snap frozen, and stored at −80° C until further use. The C-MYC/DDK tagged purified recombinant human EIF4G1, EIF4A1, and PABP proteins were purchased from OriGene (Rockville, MD).

### In vitro binding assays

Recombinant, bait GST-tagged EIF4E or 6xHis-tagged CELF1 proteins (1 μM) were initially bound to glutathione-agarose beads (ThermoFisher Scientific) or Ni^2+^ beads (GE Healthcare), respectively. Post-binding, beads were washed three times with wash buffer containing 50 mM Tris-HCl (pH 7.4) and 20% glycerol before addition of prey proteins (0.1 −10 μM). Mixtures were incubated at room temperature with rocking for 2 hours. Beads were washed three times with wash buffer containing 500 mM NaCl. The protein bound to the beads were resolved by SDS-PAGE and analyzed by both Coomassie staining and immunoblot analysis.

### Mammalian cell free in vitro translation

Wild-type or GRE deletion mutant *CRLF1* and *SNAI1* 3’ UTRs, two of the GRE-containing mRNAs, were fused downstream of the *Renilla* luciferase coding sequence in the broadly used pRL-TK CXCR4 6x reporter plasmid (Addgene, Cambridge, MA) as described before (10). T7 forward and respective 3’ UTR-specific reverse primers, listed in **Table S2**, were used to generate PCR templates for *in vitro* transcription from pRL-TK CXCR4 6x, pRL-TK *CRLF1*, pRL-TK *CRLF1* _Δ_*GRE*, pRL-TK *SNAI1*, and pRL-TK *SNAI1* _Δ_*GRE* plasmids. Capped and poly-adenylated mRNA templates were generated using the mMESSAGE mMACHINE kit and Poly(A) Tailing kit, respectively (ThermoFisher Scientific). For cap-independent translation experiments, PCR templates were generated to amplify the EMCV IRES or HCV IRES driven *Renilla* luciferase from the pFR_EMCV (10) and pFR_HCV_xb plasmid (Addgene), respectively, using primers listed in **Supplementary Table 2**. These amplicons were then used to generate the poly-adenylated mRNA template as above, except without addition of the cap analogue.

MCF10A cells were transiently transfected with shRNA targeting either *GLB1* or *CELF1* as described previously (10), and subsequently treated with TGF-beta for 72 hours. Cell-free extracts from these cells were prepared and *in vitro* translation was performed as described previously (62). Briefly, cells were lysed in freshly prepared ice-cold hypotonic lysis buffer (10 mM HEPES, pH 7.6, 10 mM potassium acetate, 0.5 mM magnesium acetate, 5 mM DTT, EDTA-free protease and phosphatase inhibitor cocktail (ThermoFisher Scientific). Where indicated extracts were treated with 4 ng/µl recombinant coxsackievirus B3 2A protease as described above. For immunodepletion of eIF4G1, extracts were processed as described above for immunoprecipitation with an anti-eIF4G1 antibody or beads alone (**Supplementary Table 2**) and the process was repeated twice.

*In vitro* translation reactions were set up in a total reaction volume of 10 µl per sample (containing 4 µl of cell-free extract, 1 µl of translation buffer (16 mM HEPES, pH 7.6, 20 mM creatine phosphate, 0.1 µg/µl creatine kinase, 0.1 mM spermidine and 100 µM amino acids (Sigma-Aldrich)), 0.4 µl of 1M potassium acetate, 0.2 µl of 100 mM magnesium acetate, 20U RNasin (Promega), and 0.04 pmol of mRNA template). Reactions were incubated out for 30 minutes at 37°C. A reaction with no added mRNA template was used as a background control. Translation read-out was performed using the dual-luciferase reporter assay system (Promega) as per the manufacturer’s protocol on a Tecan M200 multimode reader running Tecan Magellan software (Tecan). Data is presented as mean ± standard deviation (SD) of luciferase light units.

### In vitro migration and invasion assay

*In vitro* migration and invasion assays were performed using Culturex 96 Well Cell Migration and Invasion Assay kits (Trevigen, Gaithersburg, MD) as described before (10). Data obtained were used to analyze percent migration and invasion and were expressed as percent mean ± standard deviation.

### Animal studies

All mouse procedures were approved by the Institutional Animal Care and Use Committees of the Baylor College of Medicine and were performed as described before (10). Six-week-old spontaneous mutant T-cell-deficient female homozygous nude mice (NCr-*Foxn1^nu^*) (Taconic, Hudson, NY, USA) were used for the experimental metastasis experiments (n=4 in *parental* group, n=4 in wild-type *CELF1* group, and n=6 in *CELF1*^Δ^*^YAAAALP^*mutant group). To assess the experimental metastatic potential of cells, 10^6^ indicated MCF10AT1 cell variants labelled with GFP-Firefly luciferase were injected into animals via the tail vein. Mice were assessed weekly for metastasis using *in vivo* bioluminescence imaging using an IVIS Imaging System (IVIS imaging system 200, Xenogen Corporation, PerkinElmer, Waltham, MA, USA). Mice were euthanized on day 15 post tail-vein injection at which time the lungs were surgically removed and fixed using 10% neutral buffered formalin. Lungs were further subjected to hematoxylin and eosin staining and immunohistochemistry using anti-CUGBP1 antibody [3B1] (ab9549; Abcam, Cambridge, MA, USA) (1:500). Images were obtained using Axio Zoom.V16 microscope (stereo).

### Statistical analysis

Laboratory data are presented as mean ± standard error of mean (SEM) unless otherwise stated. When two groups were compared, the Student’s t test was used unless otherwise indicated and a pval < 0.05 was considered significant.

## Acknowledgments

We thank Dr. Katherine L Borden for the *FLAG-EIF4E*, and *FLAG-EIF4E* (W73A) plasmids, Dr. Yan-Hwa Wu Lee for the *GST-EIF4E* and *GST-EIF4E* (W73A) plasmids. This research was supported by NCI CA190467 to J.R.N. and CPRIT grant 170172 to J.M.R, J.R.N. is the Athena Water Breast Cancer Research Scholar of the American Cancer Society (RSG-15-088-01RMC).

## Author contributions

A.C. conceived, designed and performed experiments, wrote, reviewed and edited the manuscript; N.Z. contributed to the mouse studies, reviewed and edited the manuscript; S.D.R. contributed the MCF10A eIF4E variant cell lines, reviewed and edited the manuscript; Y.Z. performed the expression and purification of recombinant GST-tagged proteins, reviewed and edited the manuscript; R.P., N.K., E.O., S.C., L.C.R. performed experiments, data analysis, reviewed and edited manuscript; R.E.L., M.S., and J.M.R. contributed to study concept, experimental design, reviewed and edited manuscript; J.R.N. supervised the project, conceived and designed experiments, and wrote, reviewed and edited manuscript.

## Data and materials availability

All data supporting the findings of this study are either included in the manuscript or available upon request from the corresponding author.

