## Supplemental Figures and Tables for "CELF1 is an EIF4E binding protein that promotes translation of epithelial-mesenchymal transition effector mRNAs"

**This file includes:**

Online Methods

Supplementary Figures 1 - 4

Supplementary Tables 1 and 2

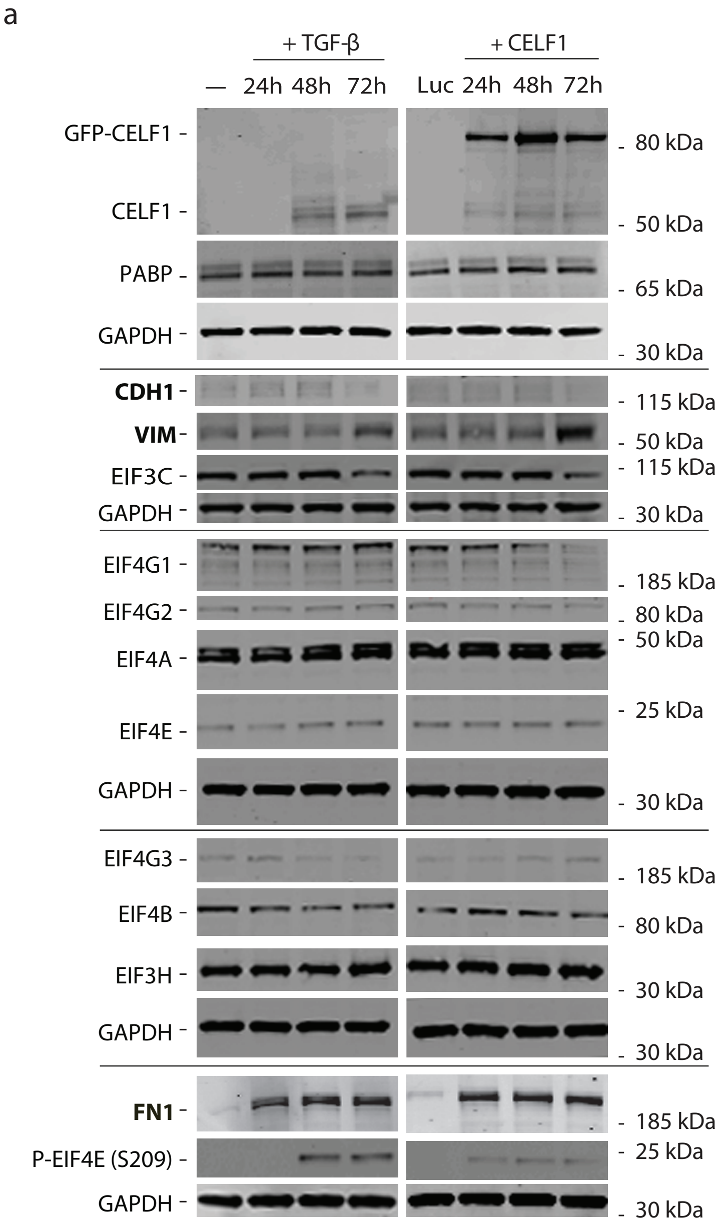


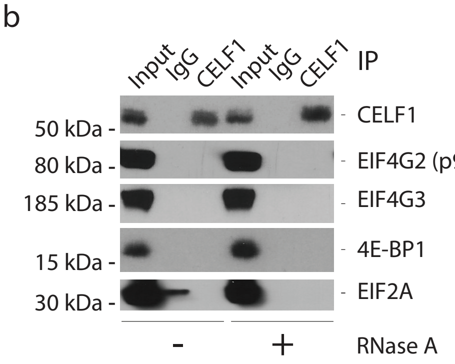


**Supplementary Figure 1. CELF1 does not interact with eIF4G2 and eIF4G3.** (**a**) Representative immunoblots of MCF10A cells induced to undergo EMT via either treatment with TGF-β (left column) or transient transfection of *GFP-CELF1* (right column). Lysates were resolved via SDS-PAGE and immunoblotted with indicated antibodies to confirm EMT induction and to profile expression and modification levels of various translational initiation factors. GAPDH = loading control; black lines separate blots derived from the same lysates but processed in parallel to enable detection of the displayed number of markers. (**b**) Immunoblots of immunoprecipitates from lysates derived from MCF10A cells treated with TGF-β for 72 hours. One half of each total immunoprecipitate was digested with RNase A prior to immunoblotting with the indicated antibodies. All results are representative of at least three individual experiments.

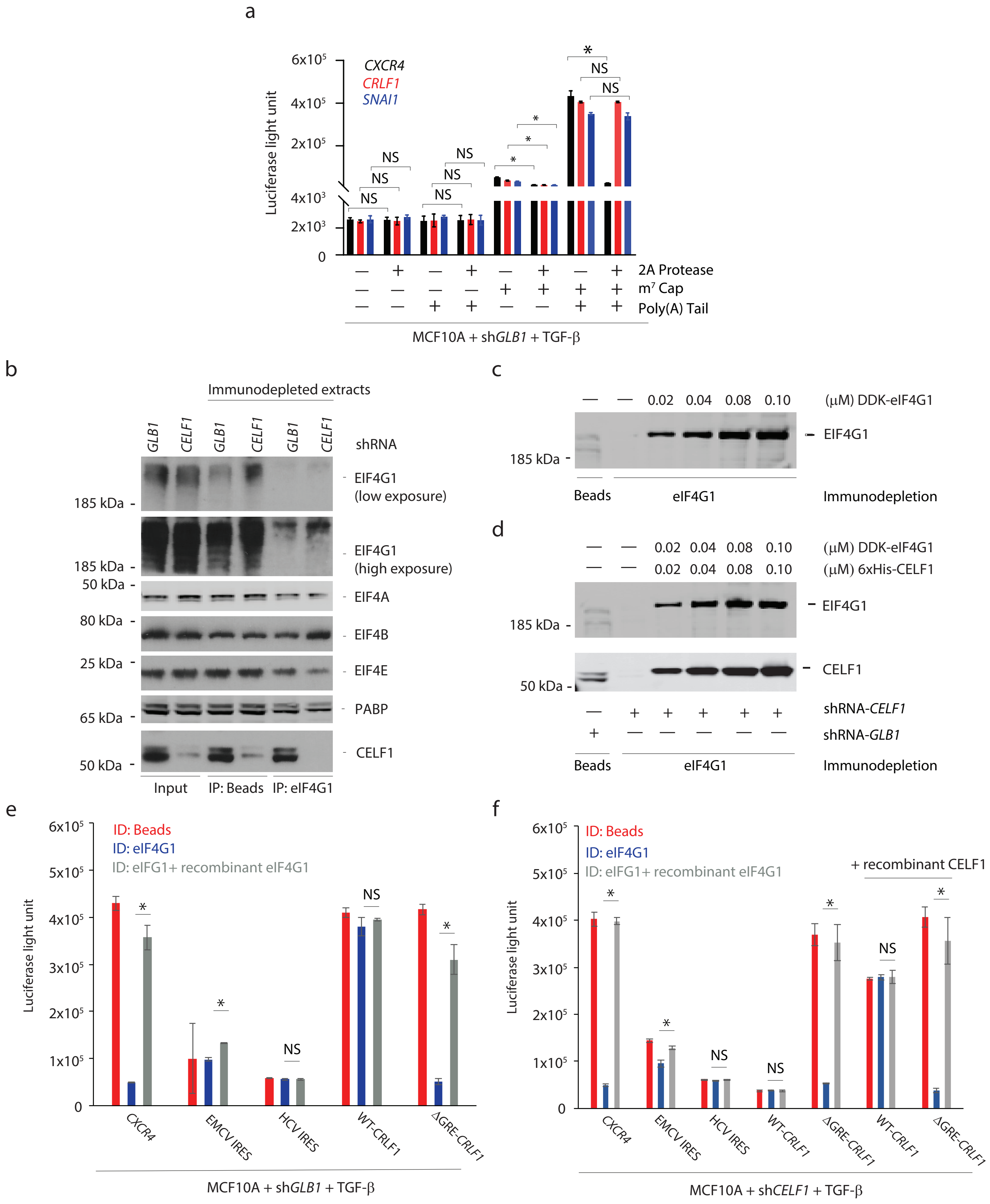


**Supplementary Figure 2. CELF1-stimulated translation of GRE-containing EMT effector mRNAs in the absence of eIF4G1 requires a m$^7$G cap and polyadenylated tail.** (**a**) Efficiency of *in vitro* translation of indicated *Renilla* luciferase reporter mRNAs in mock or 2A protease-digested cell-free extract. Presence of m$^7$G cap, polyadenylation status, and digestion status are indicated. (**b**) Representative immunoblots of eIF4G1 immunodepletion from extracts derived from MCF10A cells transiently transfected either with shRNA targeting *GLB1* or *CELF1*. Lysates were immunodepleted twice with beads (negative control), or an antibody specific to eIF4G1. Immunodepleted extracts were resolved via SDS-PAGE and immunoblotted with the indicated antibodies. (**c**) Representative immunoblot of reconstitution of eIF4G1-immunodepleted extract with purified DDK-tagged eIF4G1. (**d**) Representative immunoblot of reconstitution of eIF4G1-immunodepleted extract with purified DDK-tagged eIF4G1 and recombinant 6xHis-CELF1. (**e**, **f**) Efficiency of *in vitro* translation of reporter mRNAs as described in (a), but with mock (beads), eIF4G1 immunodepleted (ID) cell-free extract, or eIF4G1 immunodepleted extract reconstituted by addition of recombinant eIF4G1. In (a, e and f), all extracts were derived from MCF10A cells treated with TGF-β and transiently transfected with shRNAs targeting either *GLB1* (a, e) or *CELF1* (f). CXCR4 = cap-dependent control, EMCV IRES = eIF4G1-dependent but cap-independent control, HCV IRES = eIF4G1- and cap-independent control, WT = wild-type 3' UTR, ΔGRE = 3' UTR with deletion of GRE. In (a-c and d) error bars denote standard deviation. NS: not significant; *pval ≤ 0.05 (t-test). In all panels, results are representative of at least three independent experiments.

a

| PROTEIN | ORGANISM | SEQUENCE |
|---|---|---|
| CELF1 | *Homo sapiens* | GIQQYAAAALPTLYN |
| CELF1 | *Mus musculus* | GIQQYAAAALPTLYN |
| CELF1 | *Danio rerio* | GIQQYAAAALPSLYS |
| CELF1 | *Gallus gallus* | GIQQYAAAALPTLYN |
| EDEN -BP | *Xenopus laevis* | GIQQYAAAALPSLYN |
| CELF2 | *Homo sapiens* | GIQQYAAAALPTLYS |

e

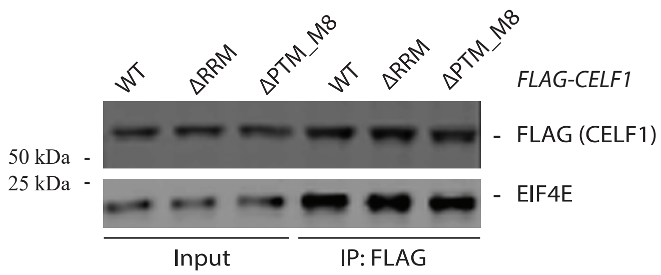


b

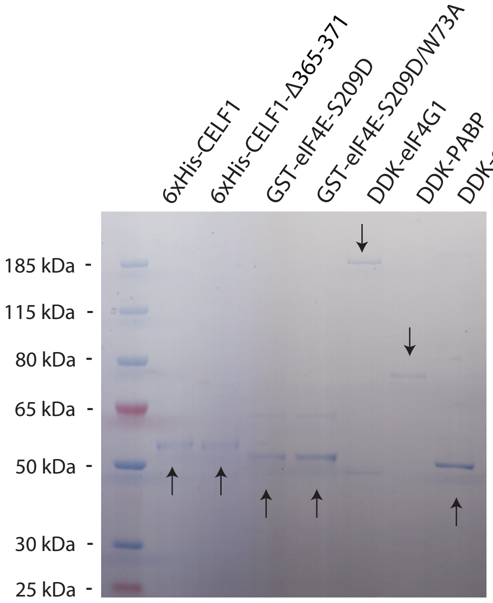


f

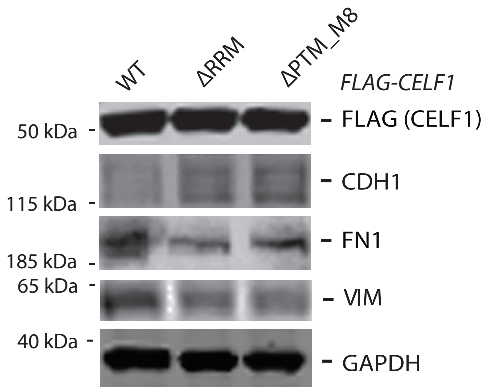


c

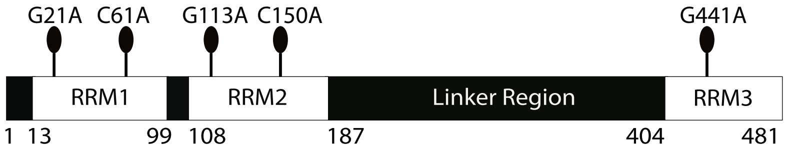


d

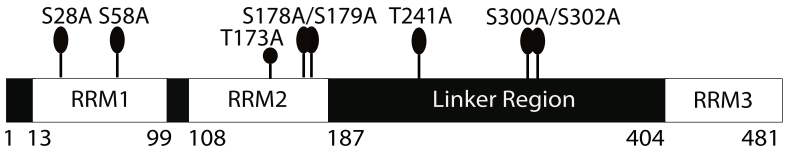


g

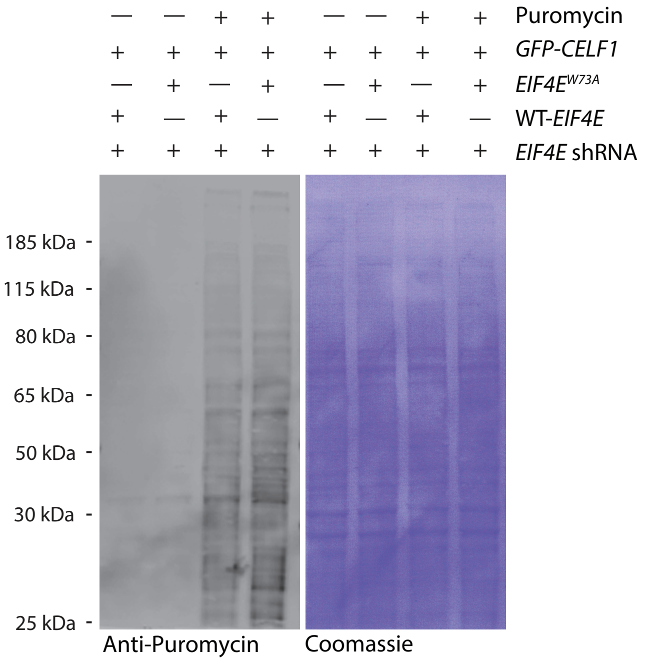


**Supplementary Figure 3. CELF1's interaction with eIF4E is independent of the former's RNA binding activity and post-translational modification (PTM).** (**a**) Multiple alignment of conserved CELF1 sequences in *H. sapiens*, *M. musculus*, *D. rerio*, *G. gallus*, and *X. laevis*, as well as *H. sapiens* CELF2. The amino acids highlighted in red in the multiple sequence alignment depict the conserved YXXXXLφ domain. (**b**) Coomassie stained gel showing relative purity of proteins used in in vitro binding assays. (**c**) Schematic representation of RNA binding mutant (ΔRRM) of CELF1. (**d**) Schematic representation of phospho-mutant form (ΔPTM-M8). (**e**) WT, ΔRRM, or ΔPTM-M8 mutant *FLAG-CELF1* constructs were transfected into MCF10A cells, which were lysed after 72 hours. Lysates were immunoprecipitated using an anti-FLAG antibody. Immunoprecipitates were resolved by SDS-PAGE and immunoblotted for FLAG-CELF1, and eIF4E. (**f**) As (e), except lysates were resolved by SDS-PAGE and immunoblotted for representative EMT markers. (**g**) Representative image of the SUnSET immunoblot used to determine the amount of puromycin-labelled peptides in MCF10A cells treated with TGF-beta for 72 hours, either expressing WT or W73A mutant phosphomimic *eIF4E* (*EIF4E-S209D*). To demonstrate the specificity in the anti-puromycin signal, non-puromycin treated cell lysates were also included. The blot was stripped and stained with Coomassie Blue R250 to confirm equal loading. All results are representative of at least three individual experiments.

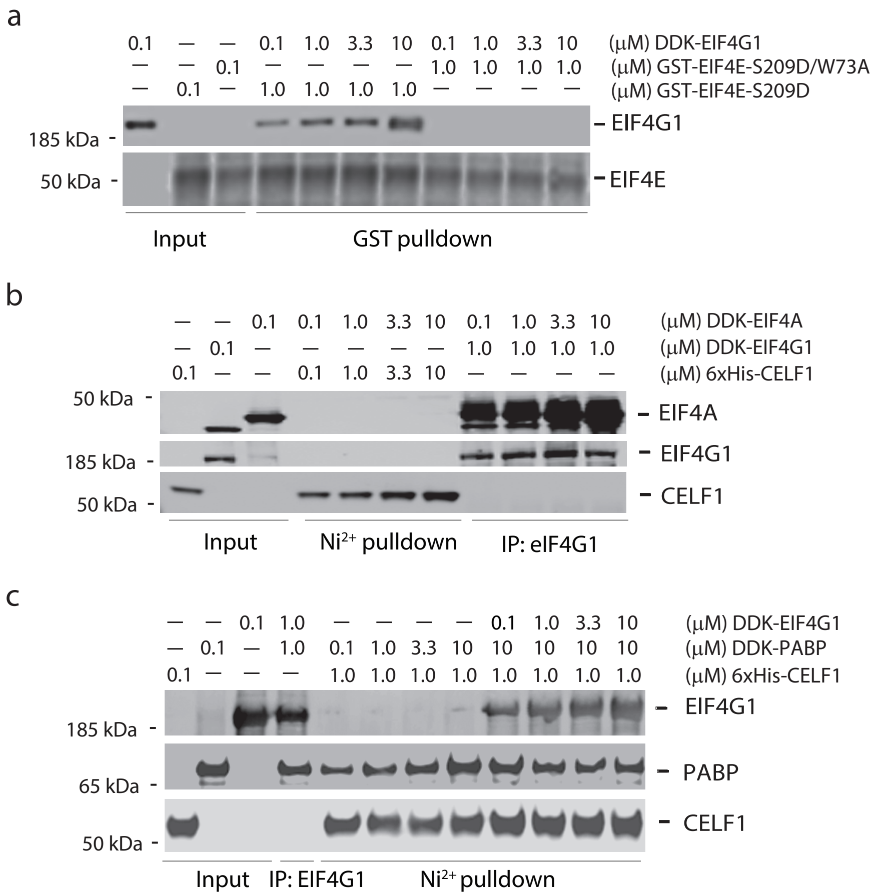


**Supplementary Figure 4. CELF1 directly binds PABP, but not eIF4A.** (**a**) Purified DDK-tagged eIF4G1 was mixed with purified GST-eIF4E-S209D or purified GST-eIF4E-S209D/W73A mutant. Complexes were pulled down with glutathione beads, washed, and immunoblotted with indicated antibodies. (**b**) Purified DDK-tagged eIF4A was mixed with purified 6xHis-CELF1 or DDK-tagged eIF4G1. Complexes were pulled down with $Ni^{2+}$ beads or immunoprecipitated (IP) using anti-eIF4G1 antibody, respectively, and immunoblotted with indicated antibodies. All results are representative of at least three individual experiments. (**c**) Purified DDK-tagged PABP was mixed with purified 6xHis-CELF1, DDK-tagged eIF4G1, or combination of 6xHis-CELF1 and DDK-tagged eIF4G1. Complexes were pulled down with $Ni^{2+}$ beads or immunoprecipitated using anti-eIF4G1 antibody, respectively, and immunoblotted with indicated antibodies. All results are representative of at least three individual experiments.

**Supplementary Table 1. List of antibodies used in the current study for immunoblot (*IB*), and immunoprecipitation (*IP*).** Concentrations used in IPs are mentioned in the relevant part of the *Materials and Methods* section). *NA*, not applicable.

| **Name of Antibody** | **Vendor** | **Catalog #** | **Dilution** |
|---|---|---|---|
| 4E-BP1 (53H11) | Cell Signaling, Beverly, MA | 9644 | 1:1000 |
| β-Actin | Abcam, Cambridge, MA | ab8227 | 1:5000 |
| β-Tubulin (D71G9) | Cell Signaling, Beverly, MA | 5568 | 1:2000 |
| CELF1 (3B1) | Abcam, Cambridge, MA | ab9549 | 1:1000 |
| E-cadherin (24E10) | Cell Signaling, Beverly, MA | 3195 | 1:1000 |
| eIF3C p110 (H-220) | Cell Signaling, Beverly, MA | 2068 | 1:1000 |
| eIF3H | Cell Signaling, Beverly, MA | 3413 | 1:1000 |
| eIF4A (C32B4) | Cell Signaling, Beverly, MA | 2013 | 1:1000 |
| eIF4B | Abcam, Cambridge, MA | ab38359 | 1:1500 |
| eIF4E | Cell Signaling, Beverly, MA | 9742 | 1:2000 |
| eIF4G1 (D6A6) | Cell Signaling, Beverly, MA | 8701 | 1:1000 |
| eIF4G1 (N-terminus) | LifeSpan Biosciences, Seattle, WA | LS-C125562 | 1:500 |
| eIF4G1 (C-terminus) (aa1230-1234) | LifeSpan Biosciences, Seattle, WA | LS-C205396 | 1:1000 |
| eIF4G2/p97 (D88P6) | Cell Signaling, Beverly, MA | 5169 | 1:1000 |
| eIF4G3 | Novus Biologicals, Littleton, CO | NBP2-16309 | 1:1500 |
| FIBRONECTIN (IST-9) | BD Biosciences, San Jose, CA | 610077 | 1:2000 |
| FOSB (5G4) | Cell Signaling, Beverly, MA | 2251 | 1:1000 |
| GAPDH | MilliporeSigma, Burlington, MA | MAB374 | 1:5000 |
| GFP | Cell Signaling, Beverly, MA | 2555 | 1:1500 |
| HA-tag (HA.C5) | Abcam, Cambridge, MA | ab18181 | 1:2000 |
| JUNB (C37F9) | Cell Signaling, Beverly, MA | 3753 | 1:1000 |
| PABP | Kind gift from Dr. Richard E Lloyd | NA | 1:2000 |

| | | | |
|---|---|---|---|
| Phospho-eIF4E (Ser209) | Cell Signaling, Beverly, MA | 9741 | 1:2000 |
| SNAIL | Abcam, Cambridge, MA | ab82846 | 1:1000 |
| SSBP2 | LifeSpan Biosciences, Seattle, WA | LS-B5585 | 1:1500 |
| VIMENTIN (RV202) | Abcam, Cambridge, MA | ab8978 | 1:1000 |

**Supplementary Table 2. Oligonucleotides used in the current study. LIC, ligation-independent cloning; RNA-IP, RNA-immunoprecipitation**

| | |
|---|---|
| **Expression Constructs** | |
| *EIF4E*-attB1 | GGGGACAAGTTTGTACAAAAAAGCAGGCTATGGCGACTGTCGAACCG |
| *EIF4E*-attB2 | GGGGACCACTTTGTACAAGAAAGCTGGGTTTAAACAACAAACCTATTTTTAGTGGTGG |
| *CELF1_LIC_F* | acctgtacttccaatccaatATGAACGGCACCCTGGAC |
| *CELF1_LIC_R* | atccgttatccacttccaatTCAGTAGGGCTTGCTGTC |

**qRT-PCR for polysome profiling and RNA-IP**

| Gene | Forward primer (5’-3’) | Reverse primer (5’-3’) |
|---|---|---|
| *ACTB* | ACCCAGCACAATGAAGATCA | ACATCTGCTGGAAGGTGGAC |
| *GAPDH* | GCGAGATCCCTCCAAAATCAA | GTTCACACCCATGACGAACAT |
| *EGR3* | GACAATCTGTACCCCGAGGA | TCCCAAGTAGGTCACGGTCT |
| *FOSB* | TCTGTCTTCGGTGGACTCCT | GAAGGAACCGGGCATTTC |
| *JUNB* | AGCTACTCCCCAGCCTCTG | GGAGGTAGCTGATGGTGGTC |
| *SNAI1* | GCGAGCTGCAGGACTCTAAT | GGACAGAGTCCCAGATGAGC |
| *SSBP2* | TAAGGGCACCATTCCTCTTG | CCCAGGAAGTCAGCCATTAC |

**qRT-PCR for *in vitro* luciferase assay**

| | |
|---|---|
| *Renilla* Forward | ATTGAATCGGACCCAGGATTC |
| *CXCR4* Reverse | CAAAGCTAACACCGGATCGC |
| *JUNB* Reverse | GTAAACGTCGAGGTGGAAGG |
| *SNAI1* Reverse | ATTCCATGGCAGTGAGAAGG |
| *SSBP2* Reverse | TGGACCCGCTTTCTTCACAAAAG |

**CELF1 putative eIF4E biding mutants**

| Gene | Forward primer (5’-3’) | Reverse primer (5’-3’) |
|---|---|---|

| | | |
|---|---|---|
| *CELF1_Δ43-49* | GGGTTTTGGCTCCTATCCACAGCACCATACTGTT | AACAGTATGGTGCTGTGGATAGGAGCCAAAACCC |
| *CELF1_Δ365-371* | TTCTGGTTGTACAGAGTTTGCTGGATACCCGAGTAG | CTACTCGGGTATCCAGCAAACTCTGTACAACCAGAA |
| *CELF1_Δ374-380* | GCGCTCCCCACTCTGCAGCAGAGTATTGGT | ACCAATACTCTGCTGCAGAGTGGGGAGCGC |

**eIF4E phosphomimic**

| Gene | Forward primer (5'-3') | Reverse primer (5'-3') |
|---|---|---|
| *EIF4E_S209D* | CAAACCTATTTTTAGTGGTGTCGCCGCTCTTAGTAGCTGTG | CACAGCTACTAAGAGCGGCGACACCACTAAAAATAGGTTTG |

***EIF4E* shRNA** TGCTGTTGACAGTGAGCGAGAGATTTGGGAGCTGAACCAATAGTGAAGCCACAGATGTATTGGTTCAGCTCCCAAATCTCGTGCCTACTGCCTCGGA

**Mammalian cell-free translation**

| | |
|---|---|
| *T7-Renilla* Forward | GAAATTAATACGACTCACTATAGGGATGACTTCGAAAGTTTATGATCCAG |
| *T7-HCV IRES-Renilla* Forward | GAAATTAATACGACTCACTATAGGGTTCTAGTGTACGTAGCCAGC |
| *T7-EMCV IRES-Renilla* Forward | GAAATTAATACGACTCACTATAGGGGCCCCTCTCCCTCCC |
| *CRLF1* Reverse | TAAAAGGACTCTTTTGGAGGG |
| *CXCR4* Reverse | CAAAGCTAACACCGGATCGC |
| *SNAI1* Reverse | GAATATCAATAAACTGTACATATAACTATA |
